# Ig-VAE: Generative Modeling of Protein Structure by Direct 3D Coordinate Generation

**DOI:** 10.1101/2020.08.07.242347

**Authors:** Raphael R. Eguchi, Christian A. Choe, Po-Ssu Huang

## Abstract

While deep learning models have seen increasing applications in protein science, few have been implemented for protein backbone generation—an important task in structure-based problems such as active site and interface design. We present a new approach to building class-specific backbones, using a variational auto-encoder to directly generate the 3D coordinates of immunoglobulins. Our model is torsion- and distance-aware, learns a high-resolution embedding of the dataset, and generates novel, high-quality structures compatible with existing design tools. We show that the Ig-VAE can be used to create a computational model of a SARS-CoV2-RBD binder via latent space sampling. We further demonstrate that the model’s generative prior is a powerful tool for guiding computational protein design, motivating a new paradigm under which backbone design is solved as constrained optimization problem in the latent space of a generative model.

## 1 Introduction

Over the past two decades, structure-based protein design has provided novel solutions to challenging problems such as enzyme catalysis, viral inhibition, *de novo* structure generation and more[1, 2, 3, 4, 5, 6, 7, 8, 9]. In the vast majority of successful design examples using the popular Rosetta framework, the protein design process consists of two steps: (1) generation of a protein backbone, and (2) design of a sequence that minimizes the folded-state energy of the generated backbone. During the first step, backbone conformations are massively sampled without amino acid identity to search for templates that can host features such as catalytic sites, or loop conformations suited to particular functions. In the second step, sequences are chosen to both sustain the folded-state structure and provide the chemical elements that perform the desired function.

While many methods have been developed to perform sequence design [10, 11], few are capable of generating backbones. Currently the majority of design templates are generated using fragment sampling in combination with expert-specified topologies of loop and secondary structure elements. Creating novel backbones for which there exist a foldable sequence remains one of the greatest challenges in protein engineering, and most engineering endeavors rely only on native structures as templates. In this study, we focus on this task of protein backbone generation, and seek to develop a method that allows us to: (1) generate novel, designable structures from the limited set of existing structural data, and (2) generate backbones that satisfy user-specified design criteria.

With recent advances in deep learning technology, machine learning tools have seen increasing applications in protein science, with deep neural networks being applied to tasks such as sequence design[11], fold recognition[12], binding site prediction[13], and structure prediction[14, 15]. Generative models, which approximate the distributions of the data they are trained on, have garnered interest as a data-driven way to create novel proteins. Unfortunately the majority of protein-generators create 1D amino acid sequences[16, 17, 18, 19] making them unsuitable for problems that require structure-based solutions such as designing protein-protein interfaces.

A major challenge in the field of 3D deep learning arises from the fact that 3D coordinates lack rotational and translational invariance, making generalizable feature learning and generation difficult. To address this challenge, our own group was the first to report a Generative Adversarial Network (GAN) that generated 64-residue peptide backbones using a distance matrix representation [20] that preserved the desired invariances. 3D coordinates were recovered using a convex optimization algorithm[20] and later, a learned coordinate recovery module[21]. Despite its novelty, the GAN method was accompanied by several difficulties. First, the generated distance matrices were not Euclidean-valid, and thus it was not possible to recover 3D coordinates that perfectly satisfied the generated distances. Second, because of the redundancy of the distance matrix representation under reflection, the quality of the torsion distributions were often degraded, leading to loss of important biochemical features, such as hydrogen-bonding, in many outputs. Ultimately, the pure distance-based representation of our GAN yielded structures that, while novel, were often chemically unrealistic and unsuitable for design [22]. Although a few other algorithms that generate contacts have been reported [23, 20, 21, 24], all of these methods require external tools to build or recover 3D coordinates.

Here, we present a new variational autoencoder (VAE)[25] architecture that is the first to perform direct 3D coordinate generation of full-atom protein backbones, circumventing the need to recover coordinates from pairwise distance constraints and avoiding the problem of distance matrix validity[26]. Our model is novel in that it provides a loss function that is rotationally and translationally invariant, while also having torsional awareness and the ability to directly generate 3D coordinates. We train our VAE using a unique loss function, which formulates coordinate generation as the solution to the joint problems of distance matrix *reconstruction* and torsion angle *inference*, both of which preserve the desired invariances. With the intention of delivering high-quality, immediately designable templates, our model performs class-specific generation of immunoglobulin (Ig) proteins, which are comprised of a two-layer beta-sandwich structure supporting variable loop regions. Immunoglobulins are highly versatile in their target-binding capabilities, serving as core components in naturally occurring antibodies and in biologics such as ScFv’s[27], nanobodies[28, 29, 30], and more.

Importantly, our model motivates a conceptually new way of solving protein design problems. Because the Ig-VAE generates coordinates directly, all of its outputs are fully differentiable through the coordinate representation. This allows us to use the Ig-VAE’s generative prior to constrain structure generation with *any* differentiable coordinate-based heuristic, such as Rosetta energy, backbone shape constraints, packing metrics, and more. By constraining and optimizing a structure in the VAE latent space, designers are able to specify any desired structural features while the model creates the rest of the molecule. As an example of this approach, we use the Ig-VAE to perform constrained loop generation, towards epitope-specific antibody design. Our technology ultimately paves the way for a novel approach to protein design in which backbone construction is solved via constrained optimization in the latent space of a generative model. In contrast to conventional methods[31, 32], we term this approach “generative design.”

## 2 Methods

### 2.1 Dataset

All training data were collected from the antibody structure database AbDb[33]. Domains that were missing residues were excluded, and sequence-redundant structures were included to allow the network to learn small backbone fluctuations. The final training set is comprised of 10768 immunoglobulins spanning 4154 non-sequence-redundant structures, including single-domain antibodies. The training set covers close to 100% of the AbDb database. Structures in the dataset vary in length from 89 to 138 residues, with most falling between 114 and 130. Since the input of our model was fixed at 128 residues (512 atoms), structures larger than 128 were center-cropped. Structures smaller than 128 were “structurally padded” by using RosettaRemodel to append dummy residues to the N and C termini. The reconstruction loss of the padded regions was down-weighted over the course of training (see Supplemental Methods). We found that the structural padding step led to slight improvements in local bond geometries at the terminal regions. All structures were idealized and relaxed under the Rosetta energy function with constraints to starting coordinates[10]. This relaxation step was done to remove any potential confounding factors resulting from various crystal structure optimization procedures.

### 2.2 Model Architecture and Training

While designing our model we sought to implement three features we feel are essential to any deep-learning-based backbone generator. First, we wanted to preserve rotational and translational invariance in our loss functions to allow for generalizable feature learning across limited datasets. Second, we wanted our model to be torsionally aware, as torsion angles play a crucial role in determining a protein’s fold as well as structural quality. Last, we wanted our model to directly generate 3D coordinates to avoid any auxiliary coordinate recovery processes, and to make structure generation end-to-end differentiable. Full differentiability allows us to gradient-optimize a latent vector while subjecting generated structures to arbitrary design constraints. Many iterations of trial-and-error yielded the model training scheme is shown in Figure 1, which satisfies all three properties.

**Figure 1:**
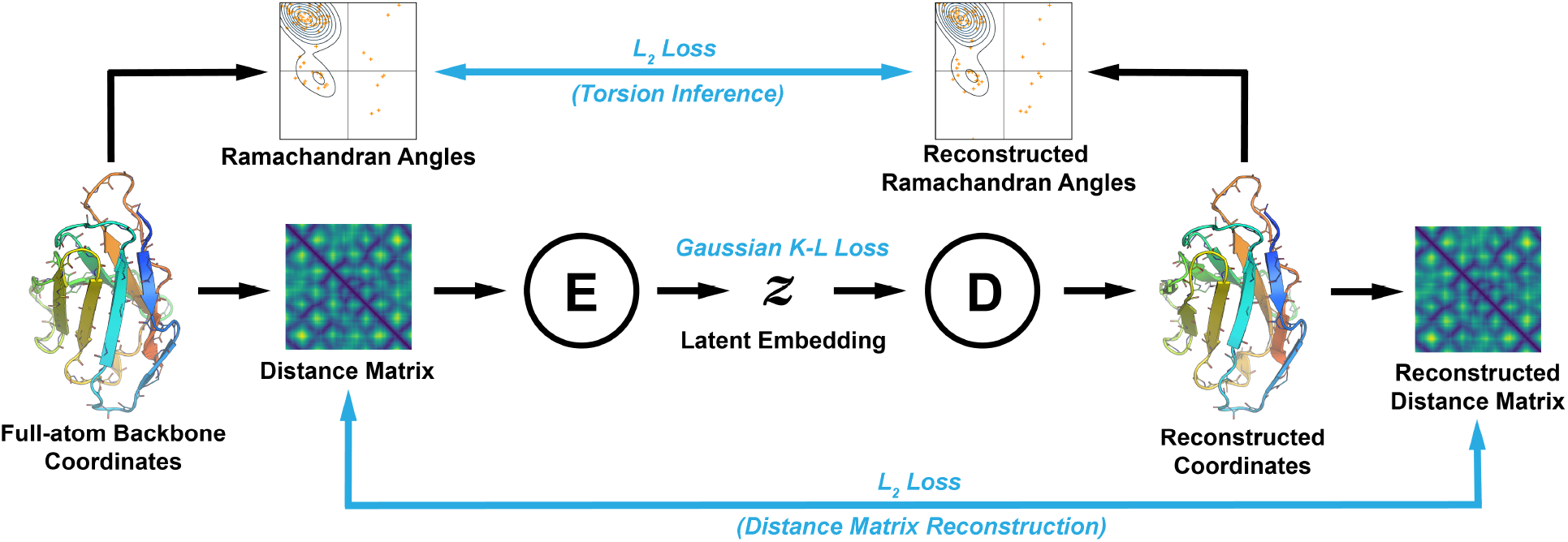
VAE Training Scheme. The flow of data is shown with black arrows, and losses are shown in blue. First, Ramachandran angles and distance matrices are computed from the full-atom backbone coordinates of a training example. The distance matrix is passed to the encoder network (E), which generates a latent embedding that is passed to the decoder network (D). The decoder directly generates coordinates in 3D space, from which the reconstructed Ramachandran angles and distance matrix are computed. Errors from both the angles and distance matrix are back-propagated through the 3D coordinate representation to the encoder and decoder. Note that both the torsion and distance matrix losses are rotationally and translationally invariant, and that the coordinates of the training example are never seen by the model. The shown data are real inputs and outputs of the VAE for the immunoglobulin chain in PDB:4YXH(L).

Like classical VAEs[25] our model minimizes a reconstruction loss and a KL-divergence loss that constrains the latent embeddings to be isotropic gaussian. During training, a distance matrix is first obtained from the coordinates of the training data. Like a conventional VAE, the encoder module compresses the distance matrix into the latent space, and the decoder is tasked with reconstructing the input data. Crucially however, the output of the decoder is *not* a distance matrix, but rather the 3D coordinates of the reconstructed structure. A reconstructed distance matrix is computed from the coordinates using a differentiable function, and loss is backpropaged from distance matrix to distance matrix, *across* the coordinate representation. Note that this reconstruction loss does not specify the absolute position of the coordinates and thus preserves the required invariances. Factoring through an explicit coordinate representation allows us to integrate a torsion loss, which we formulate as a supervised-learning objective; the network infers the correct torsion distribution from the distance matrix. This loss is computed via the coordinate representation and backpropagated through to the decoder network, and is also rotationally and translationally invariant.

We found that early in training, the torsion loss must be heavily up-weighted relative to the distance loss in order to achieve correct stereochemistry, as molecular handedness cannot be uniquely determined from pairwise distances alone. Decreasing the torsion weight later in training led to improvements in local structure quality. A detailed description of the loss weighting schedule is included in the Supplemental Methods. We note that both the distance and torsion losses are rotationally and translationally invariant, so the absolute position of the output coordinates is determined by the model itself. While coordinates of the training examples are never seen by the model directly, the model learns a natural alignment of structures along the core *β*-strands.

## 3 Results

### 3.1 Overview

To assess the utility our model, we studied the Ig-VAE’s performance on several tasks. The first of these is data reconstruction, which reflects the ability of the model to compress structural features into a low-dimensional latent space (Section 3.2). This functionality is an underlying assumption of generative sampling, which requires that the latent space capture the scope of structure variation with sufficient resolution. Next, we assessed the quality and novelty of the generated structures, characterizing the chemical validity of the samples (Section 3.3), while also evaluating the quality of interpolations between embedded structures (Section 3.4). We visualize the distribution of embeddings within the latent space to better understand its structure, and to determine if the sampling distribution is well-supported (Section 3.4). We ultimately challenge the Ig-VAE with a real design task; specifically, generation of a novel backbone with high shape-complementarity to the ACE2 epitope of the SARS-CoV2-RBD[34] (Section 3.5.1). To evaluate the general utility of our approach, we investigated whether we could leverage the model’s generative prior to perform backbone design subject to a set of local, human-specified constraints (Section 3.5.2).

### 3.2 Structure Embedding and Reconstruction

A core feature of an effective VAE is the model’s ability to embed and reconstruct data. High quality reconstructions indicate that a model is able to capture and compress structural features into a low-dimensional representation, which is a prerequisite for generation by latent space sampling. To evaluate this functionality, we reconstructed 500 randomly selected structures and compared the real and reconstructed distributions of backbone torsion angles, pairwise distances, bond lengths, and bond angles (Figure 2). The structurally padded “dummy” regions were excluded from this analysis.

**Figure 2:**
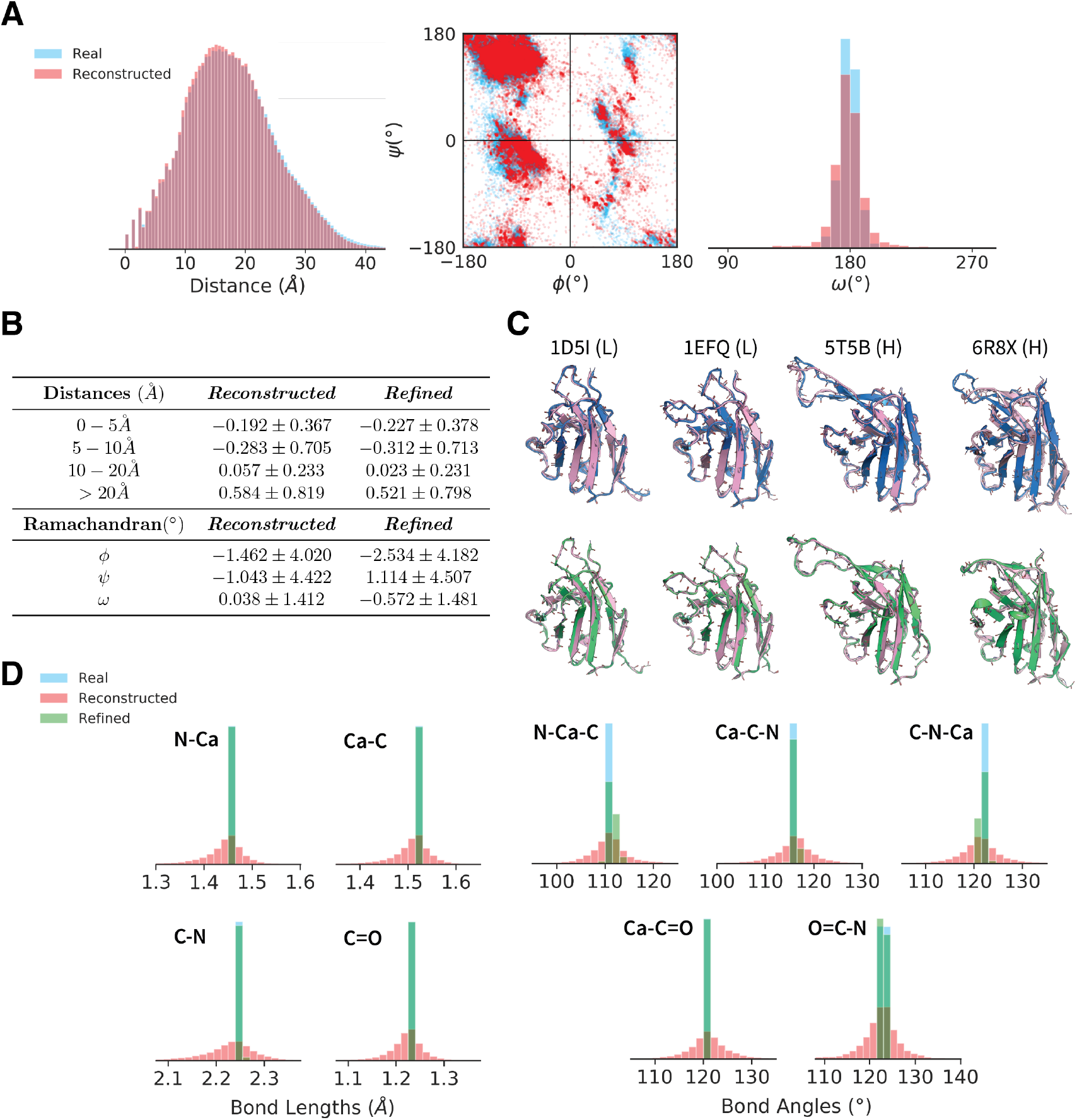
Analysis of Full-Atom Reconstructions. Reconstruction data for 500 randomly chosen, non-redundant structures in the training set. (A) Overlays of the pairwise distance and Ramachandran distributions of the real and reconstructed data. (B) A table of the reconstruction errors in pairwise distance and Ramachandran angle before and after refinement. The distance errors are reported as per-pairwise-distance error averaged over all structures in the dataset, and analogously for the angle errors. (C) Overlays of the real (blue), reconstructed (pink) and refined (green) structures. (D) Overlays of the bond length and bond angle distributions of the real, reconstructed and refined data. Overall, structures are accurately reconstructed, and errors in atom placement are small enough that they can be corrected with minimal changes to the model outputs.

The distance and torsion distributions are shown in Figure 2A, where we observe that the real and reconstructed data agree well. On average, pairwise distances smaller than 10Å tended to be reconstructed slightly smaller than the actual distances, while larger distances tended to be reconstructed slightly larger (Figure 2B, reconstructed). *ϕ* and *ψ* torsions tended within ~ 10°of the real angles, while *ω* angles tended to fall within ~ 3°. Examples of reconstructed backbones are shown in the top row of Figure 2C. These data demonstrate that the Ig-VAE accurately performs full-atom reconstructions over a range of loop conformations.

In order to use generated backbones in conjunction with existing protein design tools, it is crucial that our model produce structures with near-chemically-valid bond lengths and bond angles. Otherwise, large movements in the backbone can occur as a result of energy-based corrections during the design process, leading to the loss of model-generated features. The bond length and bond angle distributions are depicted in Figure 2D. The majority of bond length reconstructions were within ~ 0.1 Å of ideal lengths, while bond angles tended to be within ~ 10 ° of their ideal angles. We found that a constrained optimization step using the Rosetta centroid-energy function (see Supplemental Methods) could be used to effectively refine the outputs. This refinement process kept structures close to their output conformations (Figure 2C, bottom) while correcting for non-idealities in the bond lengths and bond angles (Figure 2D, green). Refinement did not improve backbone reconstruction accuracy (Figure 2B, refined), but did improve chemical validity, implying that our model outputs could be refined with Rosetta without washing-out generated structural features.

Our analysis of the reconstructions reveal that the Ig-VAE can be used to obtain high-resolution structure embeddings that are likely useful in various learning tasks on 3D protein data. These results support our later conclusion that the Ig-VAE embedding space can be leveraged to generate structures with high atomic precision, while also showing that the KL-regularization imposed on the latent embeddings does not overpower the autoencoding functionality of the VAE.

### 3.3 Structure Generation

To determine whether the Ig-VAE could generate novel, realistic Ig backbones, we sampled 500 structures from the latent space of the model and compared their feature distributions to 500 non-redundant structures from the dataset. Each generated structure was cropped based on its nearest neighbor in the training set. In Figure 3A we show overlays of the distance and torsion distributions for the real and generated structures. The generated torsions were more variable than the real torsions, with more residues falling outside the range of the training data. The real and generated distance distributions agreed well.

**Figure 3:**
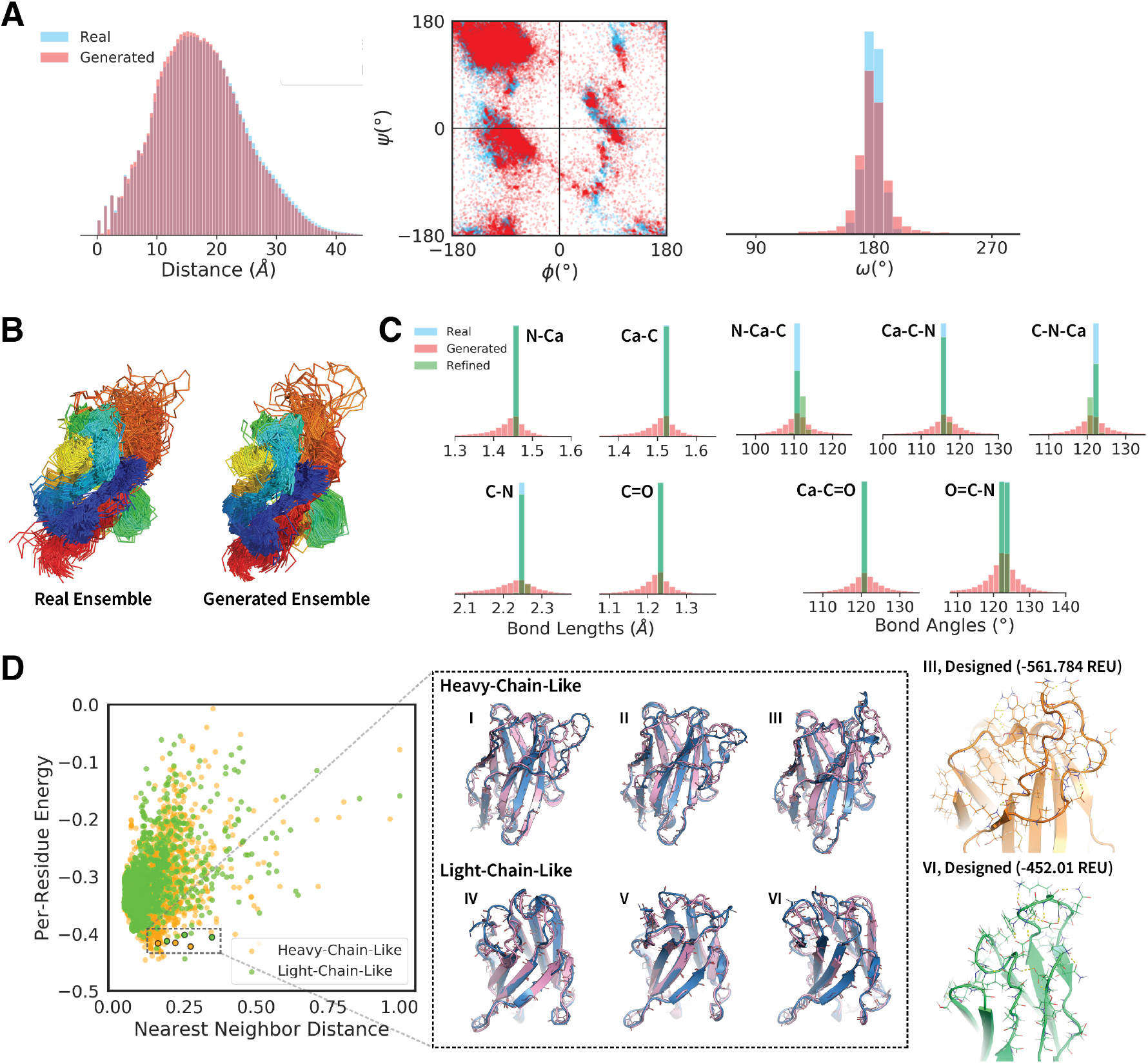
Analysis of Generative Sampling. Data for 500 of randomly selected non-redundant training samples and generated structures. **(A)** Overlays of the pairwise distance and Ramachandran distributions of the real and generated data. **(B)** A comparison of the real and generated structural ensembles. **(C)** Overlays of the bond length and bond angle distributions of the real, generated and refined data. **(D)** The left panel shows a plot of the post-refinement per-residue centroid energy against normalized nearest neighbor distance for the generated structures. The nearest neighbor distance is computed as the minimum Frobenius distance between the generated distance matrix and all distance matrices in the training set. Each point is colored based on whether the nearest neighbor is a heavy or light chain Ig. The center panel shows an overlay of the generated structures (pink) and their nearest neighbors (blue) in the training set. These six structures were selected using a combination of centroid energy, nearest neighbor distance, heavy/light classification, and manual inspection. The right panel shows sequence design results for structures **III** and **VI**. The energies in the left panel are centroid energies, while the energies in the right panel are full-atom Rosetta energies using the ref2015 score function.

Visual assessment of the backbone ensembles (Figure 3B) revealed that the two were similarly variable, suggesting that our model captures much of the structural variation found in the training set. The generated structures exhibited good chemical-bond geometries (Figure 3C, red) that were slightly noisier but comparable to those of the reconstructed backbones (Figure 2D). Once again, we found that constrained refinement using Rosetta could improve chemical bond geometries (Figure 3C, green), with minimal changes to the generated structures.

To assess both the novelty and viability of the generated examples, we evaluated structures based on two criteria: (1) post-refinement energy and (2) nearest-neighbor distance. Energies were normalized by residue-count to account for variable structure sizes. Nearest-neighbor distance was computed as a length-normalized Frobenius distance between *C_α_*-distance matrices. For notational convenience, nearest-neighbor distances were normalized between 0 and 1. We avoid the use of the classical C_*α*_-RMSD, because it is neither an alignment-free nor a length-invariant metric, and because it lacks sufficient precision to make meaningful comparisons between Ig structures.

We found that there was a positive correlation between energy and nearest-neighbor distance (Figure 3D), implying that while our model is able to generate structures that differ from any known examples, there is a concurrent degradation in quality when structures drift too far from the training data. Despite this, we found that a significant number of generated structures had novel loop shapes, achieved favorable energies, and retained Ig-specific structural features. Six of these examples are shown as raw model outputs, overlaid with their nearest neighbors in the center panel of Figure 3D. Both heavy and light-chain-like structures exhibited dynamic loop structures, and the model appears to performs well in generating both long and short loops.

To assess whether the generated loop conformations could be sustained by an amino acid sequence, we used Rosetta FastDesign[35, 36] to create sequences for the selected backbones. The outputs of two representative design trajectories are shown in the right panel of Figure 3D. The design process yielded energetically favorable sequences with loops supported by features such as hydrophobic packing, π-π stacking and hydrogen bonding. Overall these results suggest that the Ig-VAE is capable of generating novel, high-quality backbones that are chemically accurate, and that can be used in conjunction with existing design protocols to obtain biochemically realistic sequences.

### 3.4 Latent Space Analysis and Interpolation

While the results of the preceding section suggest that the Ig-VAE is able to produce novel structures, an important feature of any generative model is the ability to interpolate smoothly between examples in the latent space. In design applications this functionality allows for dense structural sampling, and modeling of transitions between distinct structural features.

A linear interpolation between two randomly selected embeddings is shown in Figure 4A. The majority of interpolated structures adopt realistic conformations, retaining characteristic backbone hydrogen bonds while transitioning smoothly between different loop conformations (Figure 4C). Structures along the interpolation trajectory were able to achieve negative post-refinement energies (Figure 4B), with the highest energy structure corresponding to the most unrealistic portion of the trajectory (Figure 4A,4B, index 20).

**Figure 4:**
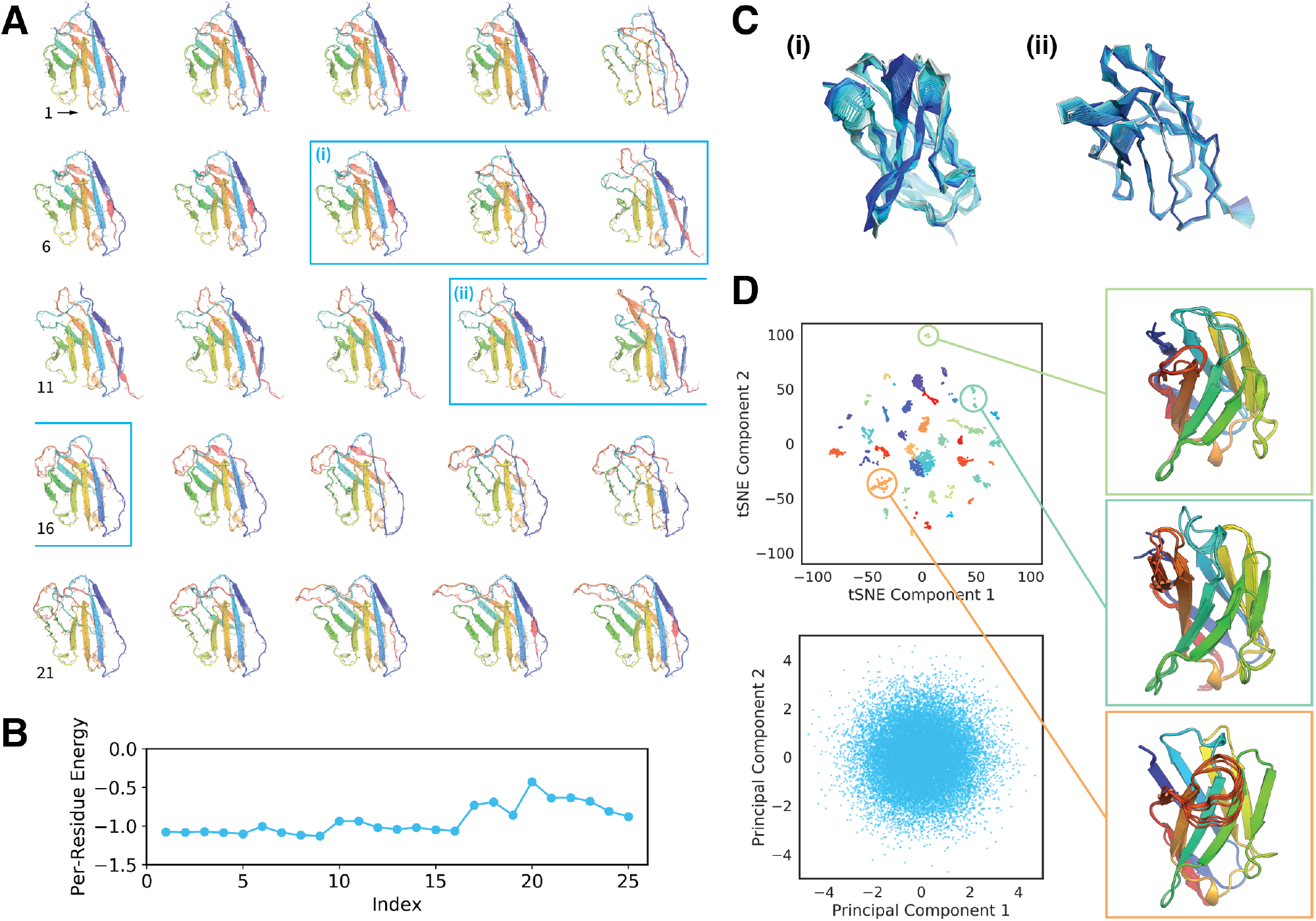
Latent Space Analysis and Interpolation. **(A)** Linear interpolation between two randomly selected embeddings. The starting and ending structures are 1TQB(H) and 5JW4(H) respectively. The backbones are unaltered, full-atom model outputs. Structures are colored by residue index in reverse rainbow order. (B) Centroid energy profiles of the structures from panel A after constrained refinement. **(C)** Higher frequency overlays of 80 sequential structures in the unrefined interpolation trajectory. The roman numerals correspond to the blue labels in panel A. A lighter shade of blue indicates an earlier structure while the darker shade indicates a later structure. Structure transitions are smooth and follow a near-continuous trajectory. **(D)** The top panel shows a tSNE dimensionality reduction of the embedding means of the 4154 non-redundant structures in the training set. Colorings correspond to k-means clusters (k=40) of the post-tSNE data, and ten structures from three clusters are visualized to the right. The bottom panel depicts a principal components reduction of the latent space, showing five sampled data points per non-redundant structure.

To better understand the structure of the embedding space, we visualized the training data embeddings (Figure 4D) using two dimensionality-reduction methods: t-distributed stochastic neighbor embedding (tSNE)[37] and principal components analysis (PCA)[38]. The top panel depicts a tSNE decomposition of the embedding means (without variance) for the 4154 non-redundant structures in the dataset. K-means clustering (k=40) revealed distinct clusters that roughly correlated with loop structure, suggesting a correspondence between latent space position and semantically meaningful features (Figure 4D, insert). In the bottom panel, we visualized sampled embeddings for the same set of structures, sampling 5 embeddings per example. PCA revealed a spherical, densely populated embedding space, suggesting that the isotropic gaussian sampling distribution is well supported. The PCA results also suggest that the KL-loss was sufficiently weighted during training.

Overall these results support the conclusions of the previous section, demonstrating that the Ig-VAE exhibits the features expected of a properly-functioning generative model, and that sampling from a gaussian prior is well motivated. The smooth interpolations agree with the observation that our model is capable of generating novel structures, which are expected to arise by sampling from interpolated regions between the various embeddings.

### 3.5 Towards Epitope-Specific Generative Design

#### 3.5.1 Computational Design of a SARS-CoV2 Binder

While antibodies are usually comprised of two Ig domains, there also exist a large number of single-domain antibodies in the form of camelids[39] and Bence Jones proteins[40]. To test the utility of our model in a real-world design problem, we challenged the Ig-VAE to generate a single-domain-binder to the ACE2 epitope of the SARS-CoV2 receptor binding domain (RBD), an epitope which is of significant interest in efforts towards resolving the 2019/20 coronavirus pandemic[34].

To do this, we sampled 5000 structures from the latent space of the Ig-VAE. To find candidates with high shape complementarity to the ACE2 epitope, we used PatchDock[41] to dock each generated structure against the RBD. To make the search sequence agnostic, both proteins were simulated as poly-valines during this step. We then selected Ig’s that bound the ACE2 epitope specifically, and used FastDesign to optimize the sequences of the binding interfaces. Two Ig’s that exhibited good shape complementarity to the ACE2 epitope and adopted unique loop conformations are shown in Figure 5A. After sequence design these candidates achieved favorable energies and complex ddG’s of −37.6 and −53.1 Rosetta Energy Units (Figure 5A, designed). Using RosettaDock[42, 43], we were able to accurately recover the designed interfaces as the energy minimum of a blind global docking trajectory, suggesting that the binders are specific to their cognate epitopes (Figure 5A, recovered, docking).

**Figure 5:**
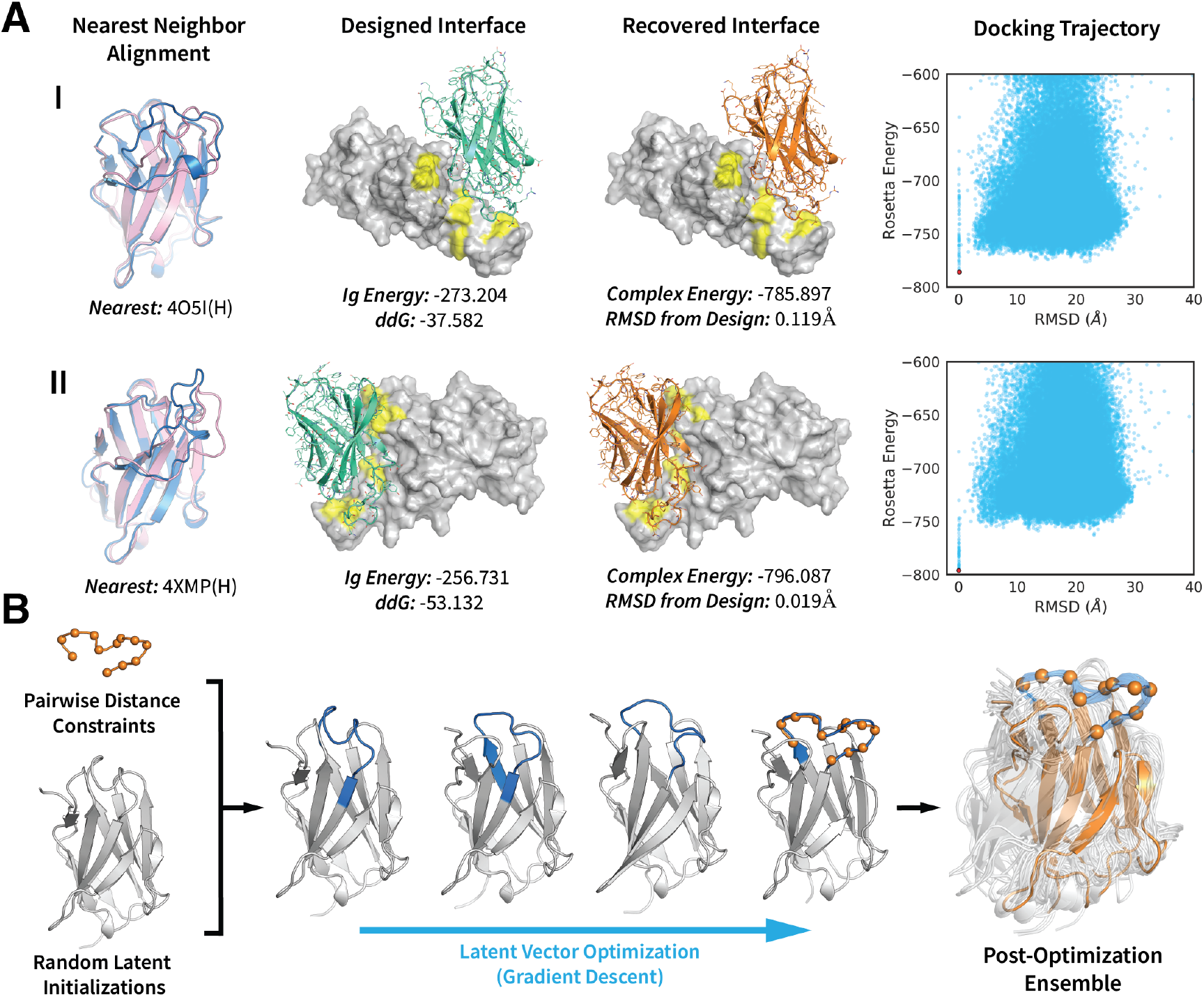
Towards Epitope-Specific Generative Design. (A) Design of two generated immunoglobulin (Ig) structures, I and II, targeting the ACE2 epitope of the SARS-CoV2 receptor binding domain (RBD). The left-most column shows an alignment of the unrefined generated structures (pink) to their respective nearest neighbors (blue) in the training set. The second column depicts the designed interfaces with the full-sequence Ig’s shown in green. The SARS-Cov2-RBD is rendered as a white surface, with the ACE2 epitope shown in yellow. The ddG’s and Rosetta energies (ref2015) of the Ig’s are shown below each complex. The third column depicts the lowest energy structures from a RosettaDock global-docking trajectory. The recovered Ig structures are shown in orange. The right-most column shows the full docking trajectory (100,000 decoys), with the lowest energy structures shown as red dots. (B) Ig-backbone design by constrained latent vector optimization. The loop constraints are taken from PDB:5X2O(L, 32-44) with the target shape shown by orange spheres. The constrained regions of the optimization trajectory are shown in blue. In the post-optimization ensemble the full 5X2O(L) backbone is shown in orange. The ensemble depicts the outputs of 62 optimization trajectories (out of 100 initialized) that successfully recovered the target loop shape.

These results demonstrate that the latent space of our generative model can be leveraged to create novel binding proteins that are otherwise unobtainable by discrete sampling of real structures. While we believe that designed proteins must be experimentally validated, our data suggest that generative models can provide compelling design candidates by computational design standards, making the method worthy of larger-scale and broader experimental testing. Our data also demonstrate the compatibility of the Ig-VAE with established design suites like Rosetta, which have conventionally relied on real proteins as templates.

#### 3.5.2 Generative Design

The functional elements of a protein are often localized to specific regions. Antibodies are one example of this, where binding is attributable to a set of surface-localized loops, as well as enzymes, which depend on the positioning of catalytic residues to form an active site. Despite this apparent simplicity, natural proteins carry evolution-optimized features that are required to host functional elements. The protein design process often seeks to mimic this organization, requiring the engineering of supporting elements centered around a desired feature. While logical, designing supportive features is almost always a difficult task, requiring large amounts of experience and manual tuning.

Motivated by these difficulties, we sought to investigate whether the generative prior of our model could be leveraged to create structures that conform to a human-specified feature without specification of other supporting features. To test this, we specified a 12-residue antibody loop shape as pairwise Ca distances. We then sampled 100 random latent initializations and applied the constraints to the generated structures. Next, we optimized the structures via gradient descent, backpropagating constraint errors to the latent vectors through the decoder network. From 100 initializations, we were able to recover the target loop shape in 62 trajectories, with the vast majority of structures retaining high quality, realistic features. We visualize one trajectory in the center panel of Figure 5B. While the middle Ig-loop (Figure 5B, blue) is being constrained, the other loops move to adopt sterically compatible conformations and the angles of the *β*-strands change to support the new loop shapes. We note that the latent-vector optimization problem is non-convex, which is why we require multiple random initializations[44, 45].

Importantly, the recovered backbones in the generated ensemble differ from the originating structure (Figure 5B, orange). These data suggest that our model can be used to create backbones that satisfy specific design constraints, while also providing a distribution of compatible supporting elements. We emphasize that this procedure is not limited to distance constraints, and can be done using any differentiable coordinate-based heuristic such as shape complementarity[46], volume constraints, Rosetta energy[47], and more. With a well-formulated loss function, which warrants a study in itself, it is possible to “mold” the loops of an antibody to a target epitope, or even the backbone of an enzyme around a substrate of choice. Our example demonstrates a novel formulation of protein design as a constrained optimization problem in the latent space of a generative model. In contrast to methods that require manual curation of each part of a protein, we term our approach “generative design,” where the requirement of human-specified heuristics constitutes the “design” element, and using a generative model to fill in the details of the structure constitutes the “generative” element.

## 4 Discussion

While a protein’s fold dictates the range of functions that it can host, both sequence features and structural features *within* a fold are crucial in dictating function. In the case of antibodies, structural variation in the loop regions plays an especially important role, often determining target compatibility and specificity [48, 49]. Although fragment sampling of existing structures has allowed us to engineer a wide range of folds [1, 2, 3], it remains a challenge to reliably generate novel Ig-loops outside the realm of known structures, making binder design for novel targets difficult.

In this study we have approached this problem by training a deep generative model that generates Ig structures with novel loop shapes, drawing only from existing structural data. We demonstrate that the structural distribution captured by our VAE can be used for backbone design, which we pose as a conditional generation problem solved via constrained optimization in the model’s latent space. Specifically, the generative prior provided by the Ig-VAE restricts backbone conformations to the regime of the Ig-fold, and the latent space search effectively generates structures conforming to user-specified features. This scheme, which we term “generative design”, is possible because our model generates 3D coordinates directly, as a result of innovations that allow for rotational and translational invariance, and torsional awareness during training.

In its current form the Ig-VAE yields a powerful tool for creating single-domain antibodies, and allows for high-throughput construction of epitope-tailored, structure-guided libraries. With such a tool, it may be possible to circumvent screening of fully randomized libraries, a large proportion of which are usually insoluble or fundamentally incompatible with the target of interest[50, 51, 52, 53]. Importantly, our approach is not specific to Ig’s, and can be applied to any fold-class well represented in structural databases such as enzymes, nucleic acid binding domains and more. Conceptually, our study offers important insight into the massive possibilities of deep generative modeling in protein design. Generative design provides a new approach that is highly flexible, allowing for the arbitrary specification of locally desired features, while leveraging the model to “fill in” the ancillary parts of the molecule. This functionality is mathematically grounded in the framework of conditional probability, making it fully data-guided while requiring minimal human intuition compared to conventional backbone design methods.

Overall, our work is of significant interest to both protein engineers and machine learning scientists as the first successful example of 3D deep generative modeling applied to protein backbone design. We speculate that our scheme will motivate further study of class-specific generative models, as well as development of differentiable loss functions that can be used, for example, to morph enzymes to host small molecule binding sites, or even mold the loops of antibodies to the surfaces of targets.

## Code Availability

Code for the working model is provided for review, and will be released on GitHub upon publication.

## Supplemental Information

Supplemental Methods, Rosetta commands, and an interpolation movie are available for download at: https://tinyurl.com/y4cao4h9.

## Author Contributions

R.R.E. and P.-S. H. conceived of the research. The codebase was written by R.R.E. Primary experiments were designed and performed by R.R.E. The docking experiments were done with assistance from C.A.C. The manuscript was written by R.R.E. and P.-S. H. with contributions from all authors.

## Acknowledgements

We thank Sergey Ovchinnikov for helpful discourse during early phases of this project, and for contributing initial code that became part of the torsion-reconstruction loss function. We thank Namrata Anand for helpful feedback and assistance. This project was supported by startup funds from the Stanford Schools of Engineering and Medicine, the Stanford ChEM-H Chemistry/Biology Interface Predoctoral Training Program and the National Institute of General Medical Sciences of the National Institutes of Health under Award Number T32GM120007. Additionally, this project was supported by the U.S. Department of Energy, Office of Science, Office of Advanced Scientific Computing Research, Scientific Discovery through Advanced Computing (SciDAC) program. This project is also based upon work supported by Google Cloud.

